# The conflict between adaptation and dispersal for maintaining biodiversity in changing environments

**DOI:** 10.1101/490722

**Authors:** Patrick L. Thompson, Emanuel A. Fronhofer

## Abstract

Dispersal and adaptation both allow species to persist in changing environments. Yet, we have limited understanding of how these processes interact to affect species persistence, especially in diverse communities where biotic interactions greatly complicate responses to environmental change. Here we use a stochastic metacommunity model to demonstrate how dispersal and adaptation to environmental change independently and interactively contribute to biodiversity maintenance. Dispersal provides spatial insurance, whereby species persist on the landscape by shifting their distributions to track favourable conditions. In contrast, adaptation allows species to persist by allowing for evolutionary rescue. But, when species both adapt and disperse, dispersal and adaptation do not combine positively to affect biodiversity maintenance, even if they do increase the persistence of individual species. This occurs because faster adapting species evolve to hold onto their initial ranges (i.e. monopolization effects), thus impeding slower adapting species from shifting their ranges and thereby causing extinctions. Importantly, these differences in adaptation speed emerge as the result of competition, which alters population sizes and colonization success. By demonstrating how dispersal and adaptation each independently and interactively contribute to the maintenance of biodiversity, we provide a framework that links the theories of spatial insurance, evolutionary rescue, and monopolization. This highlights the expectation that the maintenance of biodiversity in changing environments depends jointly on rates of dispersal and adaptation, and, critically, the interaction between these processes.

**Significance Statement:** Species can persist when the environment changes by shifting their ranges through dispersal or by adapting to the new conditions that they experience. Thus, we might expect that dispersal and adaptation in combination would increase persistence. Using a simulation model, we show that this may not be the case. Instead, species competition causes dispersal and adaptation to have conflicting contributions to biodiversity maintenance. Dispersal and adaptation each independently increase biodiversity maintenance. But when species both disperse and evolve, faster adapting species persist in their current ranges, preventing others from shifting their ranges to track environmental change. These findings highlight the need to consider ecological and evolutionary processes together, or we risk underestimating how global change will impact biodiversity.

## Introduction

When environmental conditions change, what determines whether biodiversity will be maintained? This question remains a great and pressing challenge facing ecologists and evolutionary biologists (1) because of the unprecedented magnitude of anthropogenic environmental change (2). From an ecological perspective, metacommunity theory has established that biodiversity can be maintained in changing conditions by dispersal which allows species to track favourable conditions by shifting their ranges (i.e. spatial insurance; 3). From an evolutionary perspective, genetic changes can allow populations to persist in stressful environmental conditions, which would otherwise cause extinction (i.e. evolutionary rescue; 4-6). But of course, species exist as members of ecological communities, and so their interactions with one another also impact their ability to persist (7-9). Most of our current knowledge on how species respond to environmental change is based on studies that have only considered a subset of dispersal, adaptation, and biotic interactions (1). Thus, there is pressing need for theory that integrates the processes of dispersal and evolutionary change in the context of diverse communities, if we wish to understand how biodiversity can be maintained (10).

Theory that has integrated eco-evolutionary feedbacks in competitive communities suggests that the contributions of dispersal and adaptation to the maintenance of biodiversity may be reduced when these processes interact. When dispersal rates are fast compared to rates of local adaptation, dispersal can inhibit adaptation by offering an alternative strategy for dealing with environmental change (11). In contrast, when local adaptation is fast compared to dispersal, evolutionary mediated priority effects can emerge. Also known as monopolization effects, these occur when early-arriving species adapt to the local conditions fast enough to prevent pre-adapted but later-arriving species from colonizing (12, 13). Such monopolization effects have been demonstrated to be strong drivers of community dynamics in simulated metacommunities experiencing environmental fluctuations or disturbances (14-17). Evidence of monopolization effects has also been found in a number of empirical studies (18-21). Urban et al. (10) applied this idea to changing climates, hypothesizing that monopolization effects could prevent species from tracking their climate niches. But they did not formally model this hypothesis so it is unclear how strong this effect should be and how it should depend on rates of evolution and dispersal.

Our best expectations for how dispersal and evolution should interact to affect the persistence of multispecies communities under climate change comes from a model by Norberg et al. (22). These authors found that evolution minimized extinction risks and that this was greatest when dispersal rates were low. But, in contrast with predictions from the spatial insurance hypothesis (3, 23), they found that dispersal did not reduce extinctions because it allowed competitively superior species to expand their ranges to the detriment of other species. Thus, reconciliation of these conflicting predictions is a priority for advancing our understanding of how dispersal and evolution contribute to the maintenance of biodiversity in changing conditions.

A further gap in our knowledge is on how stochasticity alters the ability of dispersal and evolution to maintain biodiversity in changing conditions. The predictions of Norberg et al. (22) and models of spatial insurance effects have all been deterministic (3, 23). Yet, stochasticity is key to the emergence of monopolization effects in disturbed or fluctuating environments (14-17). Thus, although monopolization effects are hypothesized to prevent species from tracking environmental change through dispersal (10), we do not have good expectations of how this will depend on the rates of dispersal or levels of genetic variation. Nor do we understand the consequences of monopolization effects for the maintenance of entire assemblages of interacting species as the environment changes.

Here we use a stochastic individual-based metacommunity model to test the hypotheses that 1) when species disperse, but do not evolve their environmental optimum, biodiversity is preserved through spatial insurance, 2) when species evolve their environmental optimum, but do not disperse, biodiversity is preserved through evolutionary rescue, and 3) when species both evolve their environmental optimum and disperse, biodiversity is preserved through a combination of spatial insurance and evolutionary rescue, but monopolization effects lead to the loss of biodiversity. Thus, we expect that fast adapting species will monopolize local habitats and impede species sorting, and thus threaten the persistence of slower adapting species. Furthermore, we expect that dispersal will only provide spatial insurance in regions of the metacommunity that contain analogue environments—that is, post change conditions that fall within the initial range of environmental conditions present in the metacommunity (24). In contrast, evolutionary rescue should be possible under both analogue and non-analogue conditions (22). Monopolization effects should occur whenever evolution and dispersal combine (13).

## Results and Discussion

Overall, we find that dispersal and adaptation both independently allow species to persist during environmental change (Fig. 1a, b), consistent with the spatial insurance and evolutionary rescue hypotheses and recent findings (3, 4, 23, 25). However, in combination, dispersal and adaptive evolution can facilitate monopolization effects, which results in fewer species persisting than when either process operates on its own (Fig. 1c). Dispersal, in the absence of evolution, increases the proportion of species that persist (Fig. 2a), by providing spatial insurance (3), whereby species track their environmental optimum through shifting their ranges. Spatial insurance is only possible when local conditions after environmental change fall within the range of initial conditions in the metacommunity (i.e., analogue environments). We see that the positive effect of dispersal on species persistence increases with dispersal rate, only decreasing slightly at the very highest dispersal rates, when source-sink dynamics become so strong as to detrimentally impact source population sizes. Evolution, with no dispersal, also results in an increased proportion of species that persist, by allowing for evolutionary rescue (4) as species change their environmental optima through adaptation to new environmental conditions (Fig. 1b, 2).

**Figure 1.**
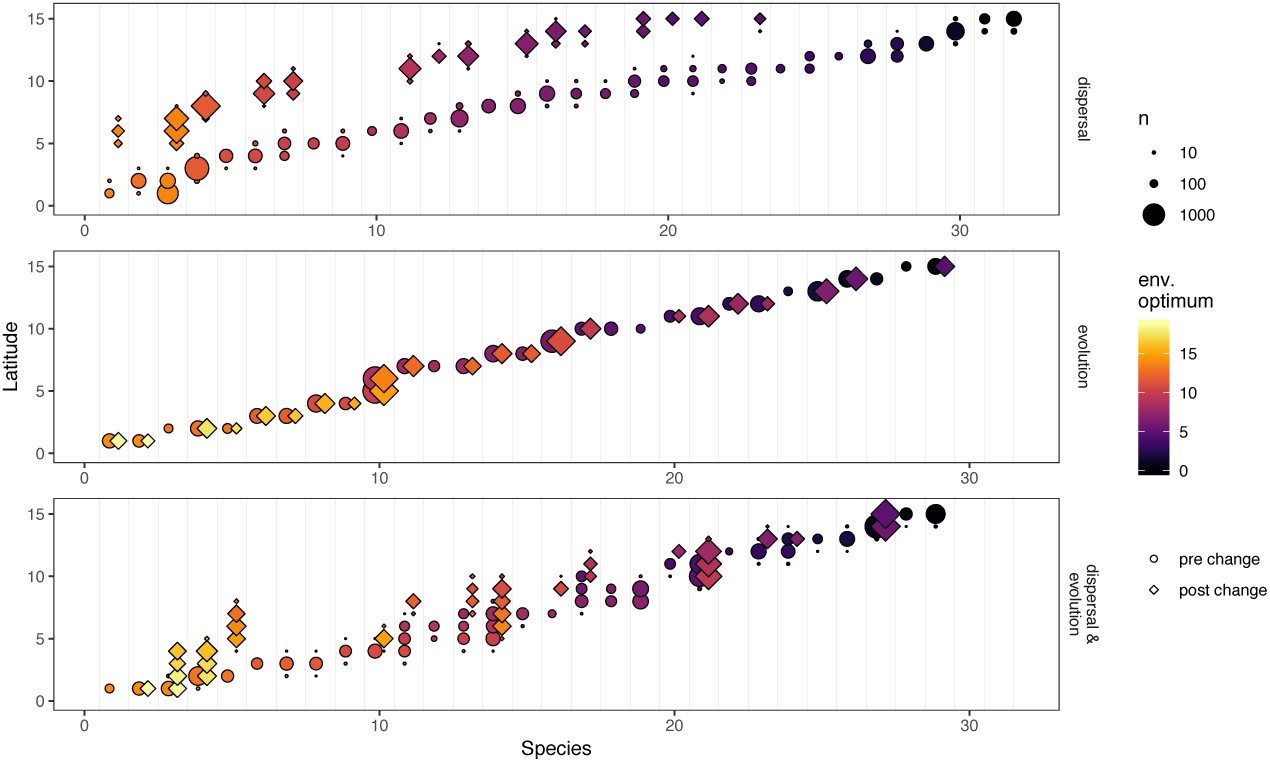
Illustration of how dispersal and evolution of environmental optimum, in isolation (a,b, respectively) and in combination (c), affect how species respond to environmental change. Only one half of the landscape ring is shown (see Fig. S1 for full landscape). Species are arranged on the x-axis by their pre-change latitude. Circles and diamonds are paired for each species and indicate the latitude and size of each population prior to and after environmental change, respectively. Species with circles but not diamonds failed to persist. The colour shows the mean environmental optimum in each population. Panel a shows a scenario where dispersal is intermediate (0.01) and the adaptive potential is zero; here, species respond to environmental change by shifting to higher latitudes to maintain the match between their phenotype and their local environmental conditions. Panel b shows a scenario where dispersal is 0 and adaptive potential is intermediate (1.08 × 10^−4^); here species respond to environmental change through adaptation (change in colour), with no change in latitude. Panel c shows a scenario where dispersal (0.01) and adaptive potential (1.08 × 10^−4^) are both intermediate; here species respond through a combination of shifting latitude and through adaptation. Results shown are from one representative simulation run with standard parameter values (Table S1). To explore additional combinations of dispersal and adaptive potential in a Shiny app visit – https://shiney.zoology.ubc.ca/pthompson/Meta_eco_evo_shiny/.

**Figure 2.**
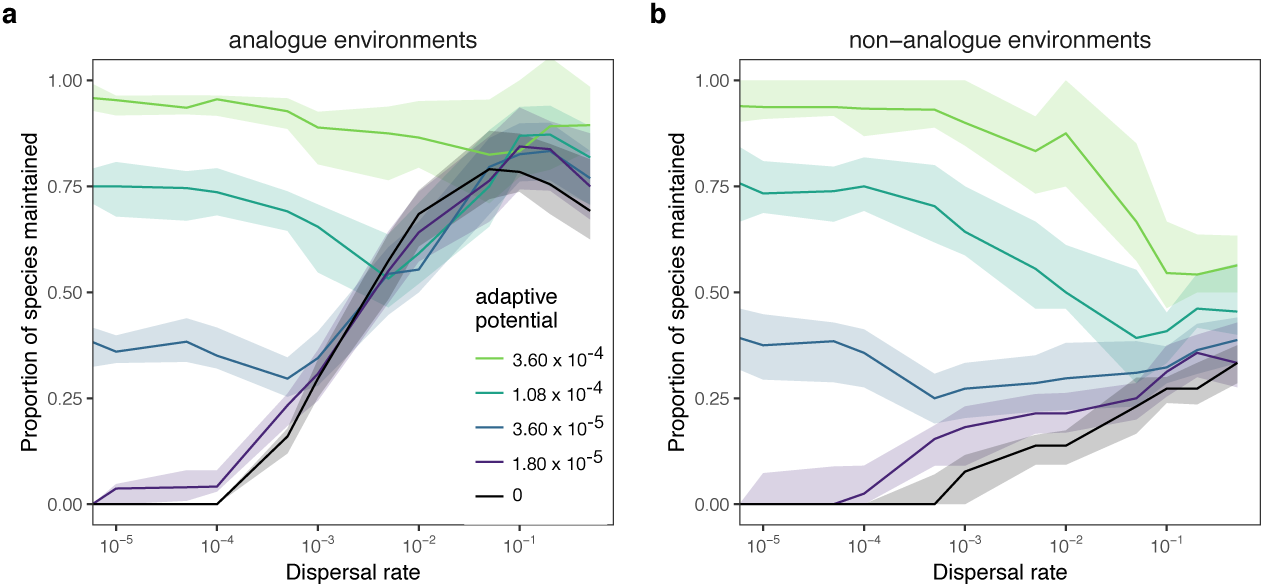
The proportion of species that are maintained following environmental change depending on dispersal and adaptive potential (colour). The proportion of species maintained was calculated as the number of species that were present in the region (a – analogue or b – non-analogue) after environmental change, divided by the number of species that were present before. Therefore, species that were only present in analogue patches would not be included in the non-analogue diversity. The lines show the median value across 50 replicate simulations with standard parameter values (Table S1) and the bands show the interquartile range. This figure shows patterns for regional scale diversity. Local scale patterns are shown in Fig. S9.

When species both disperse and evolve their environmental optima, we find that these processes generally conflict in their contributions to the maintenance of biodiversity in changing conditions (Fig. 2). That is, we see that almost all combinations of dispersal and adaptive potential result in a reduction in the proportion of species that persist, compared to the persistence that is possible with one of the two processes acting in isolation. First focusing on analogue environments, high adaptive potential (≥1.08 × 10^−4^) without dispersal is sufficient to allow the majority of species to persist (Fig. 2a). When species also disperse, the number of species that persist at these levels of adaptive potential is reduced (Fig. S2a, upper-left quadrant). Such high levels of adaptive potential may correspond to a wide array of organisms, ranging from short lived organisms with large population sizes such as plankton and other microbes, where evolutionary rescue has been commonly witnessed (6), to vertebrates with high levels of standing genetic variation (e.g. adaptation of cold tolerance in sticklebacks (26) and local adaptation across and elevational gradient in parrotbeaks (27)).

In contrast, adaptation can either increase or decrease persistence with a given dispersal rate (Fig. 2a). In particular, we see that all but the very highest levels of adaptive potential reduce species persistence when dispersal rates are intermediate (Fig. S2a, lower-left quadrant). For example, when dispersal is 0.01, increasing adaptive potential from 0 to 1.08 × 10^−4^ reduces the number of species that persist regionally from 0.74 to 0.59 (Fig. 2a). This is an intermediate level of dispersal, which has been demonstrated to maximize species sorting and environmental tracking in other metacommunity models (3, 28) and which is expected by theory to lead to classical metapopulation dynamics (29). Likewise, intermediate dispersal rates reduce the effectiveness of intermediate levels of adaptive potential in preserving species; with an adaptive potential of 1.08 × 10^−4^, the proportion of species that persist decreases from 0.75 when dispersal is 0 to 0.59 when dispersal is 0.01. Such intermediate levels of dispersal and adaptive potential likely correspond to species that are already shifting their ranges or showing increased tolerance to warming conditions, but where rates may not be high enough to ensure long term persistence (30-32). It is only with high rates of dispersal (≥0.1; e.g. as reported in metapopulations of the butterfly *Melitaea cinxia* (33)) and intermediate adaptive potential (3.6 × 10^−5^ or 1.08 ×10^−4^) that persistence is greater when evolution and dispersal combine, compared to what is possible with either process in isolation (Fig. S2a, upper-right quadrant).

This antagonistic interaction between dispersal and adaptive potential occurs as the result of monopolization effects (13), whereby species that are able to adapt faster can remain in place as the environment changes, making it harder for slower adapting species to persist by shifting their distributions. Differences in the speed of adaptation are an emergent property of our stochastic model: the first species in which an advantageous mutation arises and spreads will increase in number which increases the population scaled adaptive potential and reduces genetic drift, leading to a positive feedback loop in terms of potential for adaptation. In contrast, species that by chance don’t receive adaptive mutations early on will suffer from maladaptation and decrease in size, leading to a negative feedback loop (34, 35). As species are lost from the landscape, dispersal allows the remaining species to expand their ranges (Fig. 3a) taking advantage of reduced interspecific competition. Thus, we see the greatest range expansions when dispersal rates are intermediate and low adaptive potential results in low species persistence.

**Figure 3.**
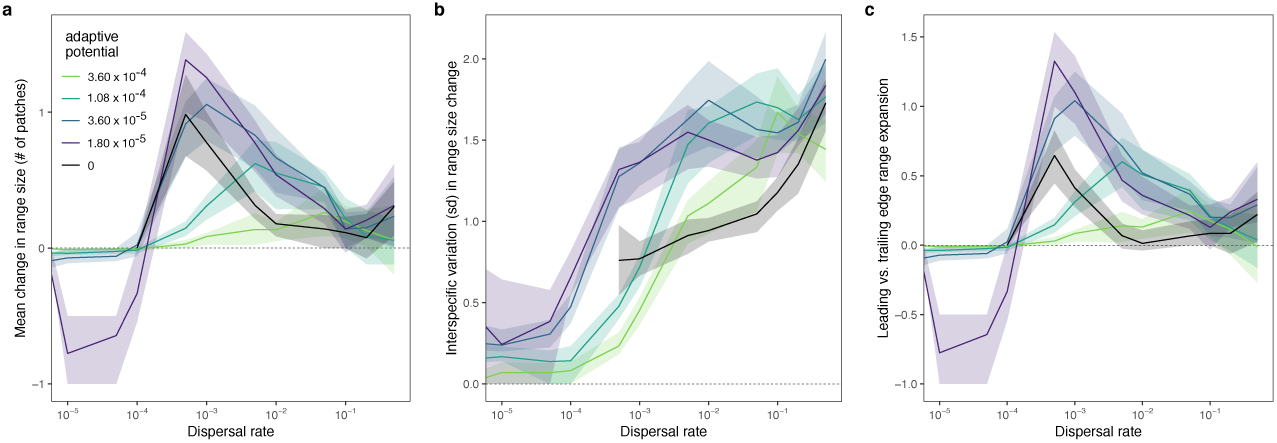
Change in the average number of habitat patches occupied (range size) by the species that persist during environmental change (a), the interspecific variation in range size change, excluding species that go extinct (b), and the leading vs. trailing edge asymmetry of the range expansion (c), depending on dispersal and adaptive potential (colour). Positive (negative) values of range change asymmetry indicate that the centroid of the range shifted towards warmer (colder) conditions, relative to the mean environmental optima of the species. The lines show the median value across 50 replicate simulations with standard parameter values (Table S1) and the bands show the interquartile range. Analogue and non-analogue regions are included together in these estimates. The lines for adaptive potential = 0 do not extend to the lowest dispersal rates because all species went extinct during environmental change.

A signature of the monopolization effect is interspecific variation in the degree to which species expand their ranges (Fig. 1c, 3b). When dispersal or adaptation occur in isolation, we see relatively little variation in the number of patches that the remaining species occupy. This variation increases with increasing rates of dispersal (Fig. 3b), but intermediate adaptive potential results in even greater variation in range size change at any given dispersal rate because they facilitate monopolization. Monopolization occurs when faster evolving species expand their distributions as conditions change (Fig. 1c), with adaptation allowing them to remain in place on their trailing (e.g., warm) edge, and dispersal allowing them to expand their ranges on their leading (e.g., cold) edge (Fig. 3c). This monopolization of the landscape by a few species causes the other species to go extinct or to be restricted to only a few patches. Consequently, we see the greatest difference in interspecific variation in range size compared to the no adaptation scenarios when adaptive potential and dispersal rates are intermediate (Fig. 3b). These scenarios correspond to the region of parameter space where dispersal and adaptation potential have the greatest conflict in the contribution to species persistence (Fig. 2a).

An additional signature of the monopolization effect is that we see leading range boundaries shifting faster than trailing range boundaries, with intermediate adaptive potential and dispersal (Fig. 1c, 3c). Species hold onto their trailing edge (i.e., warm boundary) by adapting as the environment changes, but the range expands on the leading edge (i.e., cold boundary) through dispersal. This pattern corresponds with the observation that species ranges tend to shift faster at leading edges compared to trailing edges in response to climate change (36-38) (but see 39).

In non-analogue conditions, that is, regions where the final environmental conditions exceed those present in the initial metacommunity, adaptive potential is equally effective at preserving species diversity, but dispersal is far less effective (Fig. 2b). Furthermore, dispersal reduces the effectiveness of adaptive potential for preserving species richness (Fig. 2b) because it reduces rates of adaptation through gene swamping (Fig. S3). The exception occurs when adaptive potential is zero. In this case, persistence in non-analogue environments increases with dispersal, but this is simply due to the fact that high dispersal facilitates source-sink dynamics so that species are able to persist in non-analogue patches that are in close proximity to analogue patches, where population growth rates are positive. This follows the conclusions of Norberg et al. (22), that biodiversity maintenance will depend most on evolutionary processes in regions of the planet where climate change creates non-analogue conditions (40), and that dispersal has the potential to reduce evolutionary rescue in these regions. Maintaining biodiversity in regions that have no current climate analogue will likely be a major challenge. However, as non-analogue climatic regions are expected to cover a minority of the globe (40), we have elected to focus mostly on analogue regions here.

### Biotic interactions and persistence under change

We find that diversity losses result from asymmetric responses of the species to environmental change that are driven by competition between species. Without interspecific interactions (Fig. S4) all species are able to persist, except in the limiting cases when dispersal or adaptive potential is so low that persistence is not possible. Competition results in asymmetric responses to environmental change through two mechanisms. First, it causes unequal reductions in equilibrium abundances. Species with larger populations are more likely to persist because they tend to contain more genetic variation, making adaptation faster (4) and because they produce a greater number of dispersing individuals, making range shifts more likely (23). Second, competition alters the response of species to local environmental conditions and environmental change (7, 9). Thus, competition results in interspecific differences in the ability of species to colonize new habitats in order to track environmental change through species sorting. These differences in colonization abiolity occur because the resident community acts as a filter, allowing some species to colonize, while repelling others, even if colonizing species are equally adapted to the local environment.

### The interaction between dispersal and biotic interactions

Dispersal acts to maintain biodiversity as the environment changes by providing spatial insurance, whereby species shift their distributions to ensure that they are locally adapted. Without interspecific competition, relatively low rates of dispersal (dispersal ≥ 0.001) are sufficient to allow almost all species to persist as the environment changes (Fig. S4). However, with interspecific competition, higher rates of dispersal are required for maintaining biodiversity (Fig. 2a). This occurs because high rates of dispersal generate source-sink dynamics, which counteract the effects of biotic interactions in two ways. First, dispersal spreads populations out across more patches (Fig. S5a), reducing the strength of intraspecific competition. Reduced intraspecific competition increases regional population sizes (Fig. S4a; 41), making species less prone to extinction. Second, dispersal allows species to maintain sink populations in marginal conditions, where abiotic conditions are suboptimal, but persistence would be possible without competition. Dispersal overcomes the resistance of the biotic community by providing a constant flow of immigrants. Then, if the environment changes, these populations are already in place and pre-adapted to the new conditions which makes them new potential source populations. We see that the positive effect of dispersal on regional species richness increases with dispersal (Fig 2a), only decreasing slightly at the very highest dispersal rates, when source-sink dynamics become so strong as to detrimentally impact source population sizes (see also 23). This beneficial effect of source-sink dynamics is somewhat unexpected, as we tend to consider sink populations to be a drain on the overall metapopulation. However, our results suggest that source-sink dynamics may play a key role in providing spatial insurance in changing environments.

### The interaction between evolution and biotic interactions

In contrast, adaptation acts to maintain biodiversity by facilitating evolutionary rescue. Without interspecific competition, moderate levels of adaptive potential are sufficient for maintaining all species (Fig. S4). With competition, higher levels are required (Fig. 2a) because competition reduces population sizes, increasing drift and resulting in fewer mutations (42). Of course, biotic interactions could also increase the potential for evolutionary rescue if these interactions select against maladapted individuals (42, 43). However, in our case this selective boost does not occur. Rather, competition acts to slow down rates of evolutionary rescue by reducing population sizes.

### Dispersal, evolution, and biotic interactions

When dispersal, adaptation, and competition are combined, monopolization effects are possible, which reduce the ability of spatial insurance and evolutionary rescue to preserve regional biodiversity. Monopolization effects occur when adaptive potential and dispersal rates are intermediate (Fig. 2a) and are only possible because of interspecific competition. Competition leads to interspecific differences in population sizes. Species with larger population sizes are more likely to adapt to the changing conditions and they have a higher probability of colonizing new habitats because they produce more dispersers. These species begin to occupy more space on the landscape, to the detriment of their competitors, which can lead to extinctions and the loss of diversity (Fig. 2a). Previous consideration of the monopolization hypothesis has mostly focused on static or fluctuating environments, with studies highlighting the potential for early colonizers of a habitat to become locally adapted and then repel later arriving colonists (12-14, 17, 44). However, our results support the hypothesis of Urban et al. (10) that monopolization effects may impede species sorting under climate change.

### General discussion

Our findings allow us to link the concepts of spatial insurance, evolutionary rescue, and monopolization within a common theoretical framework (Fig. 4). These processes have each been demonstrated to independently mediate the persistence of species in changing environments (3, 4, 15), but how they relate and interact with each other has not been previously shown. Critically, our study explores a range of dispersal rates and levels of adaptive potential that covers the full range of species persistence outcomes: from full extinction to levels where further increases in dispersal or adaptive potential would not increase persistence. This distinguishes our study from that of Norberg et al. (22) who demonstrated clear benefits of evolutionary responses to climate warming, but not spatial insurance. Rather, they found negative effects of dispersal on species persistence, thus precluding any potential for monopolization to reduce the effectiveness of spatial insurance. The lack of a positive effect of dispersal on species persistence is surprising, and contrasts with our findings and those of previous studies that have demonstrated spatial insurance effects (3, 23). However, it seems likely that the negative effect of dispersal in Norberg et al. (22) is due to the fact that environmental change appears to have had a relatively low impact on population performance relative to the effects of competition in their model. Without competition, all species persist and are present in all patches in the landscape, before and after environmental change. Thus, species do not need to shift their distributions in order to persist, but they can be driven extinct by competition with another species that does track its environmental optima through dispersal. Such competitive exclusion is likely in their model because of the assumption of equal inter- and intraspecific competition, which precludes stable local coexistence. Thus, the contrast between models highlights the fact that the outcome of dispersal is likely to depend on the balance in strength between the direct effects of environmental conditions and the density dependent effects of competition on the response of species to environmental change. Dispersal should have little benefit for species persistence if species distributions and the composition of species are predominantly structured by biotic interactions. But we predict that the contribution of dispersal to the maintenance of biodiversity should increase with the degree to which environmental change impacts population performance.

**Figure 4.**
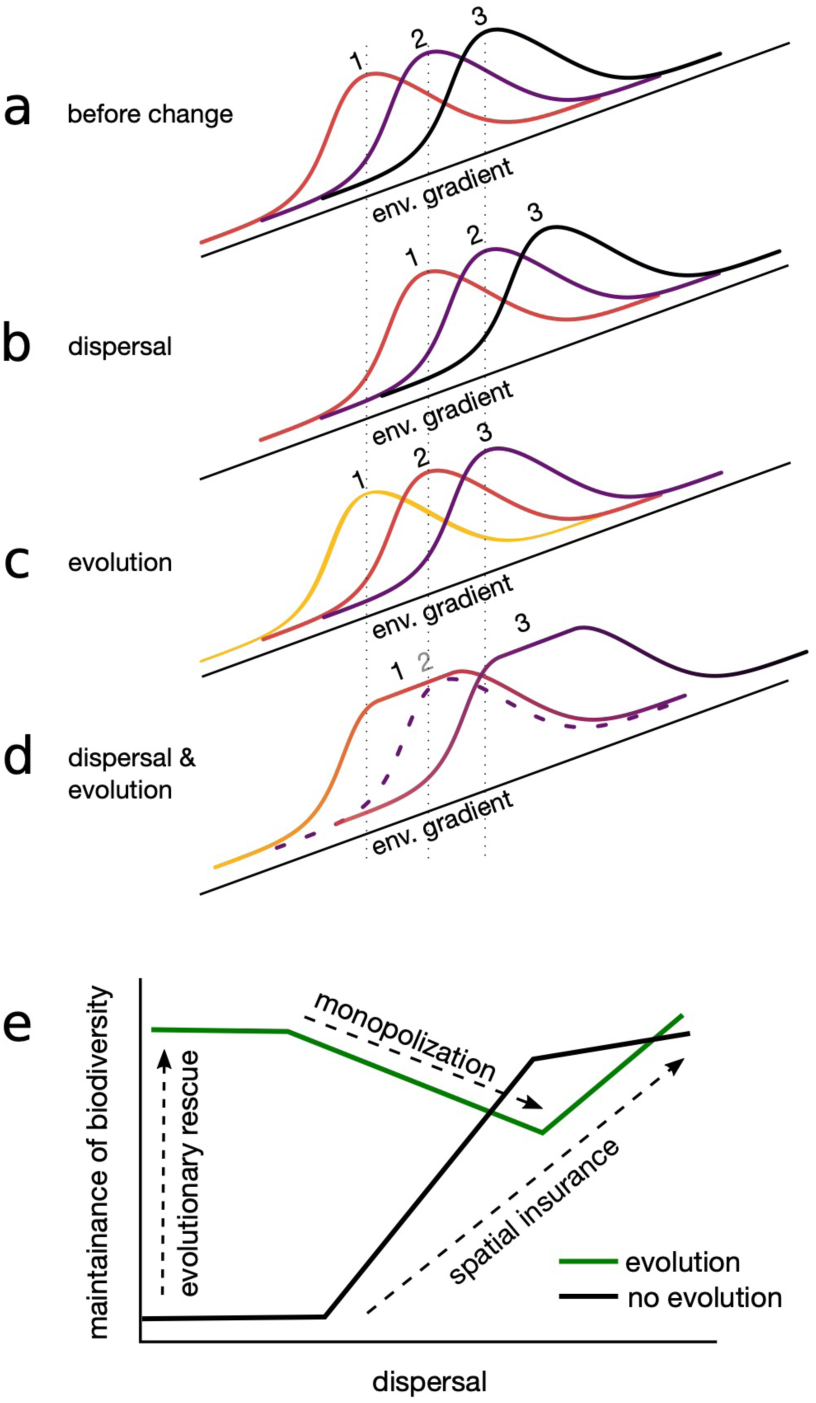
Conceptual illustration of how dispersal and the evolution of environmental optima independently and interactively act to maintain biodiversity in changing environmental conditions. Panel a shows the distribution and abundance of three species spanning a climate gradient (e.g., warm to cold). Mean position shown by vertical dashed lines. Each species is locally adapted to a different part of this gradient as indicated by the warmth of the color of the curves. Panels b, c, and d show hypothetical scenarios after the environment has changed. In panel b, with dispersal but no evolution of environmental optima, the species persist by shifting upwards (spatial insurance). In panel c, with evolution of environmental optima but no dispersal, the species persist by adapting to the changed conditions (change in color – evolutionary rescue), but do not shift their ranges. In panel d, with both dispersal and evolution, the two outer species evolve faster than the middle species, holding onto their initial trailing edge through adaptation but expanding on their leading edge through dispersal (monopolization). By monopolizing the landscape, they drive the middle species extinct (dashed curve). Panel e shows how dispersal and the evolution of environmental optima each allow for species persistence via spatial insurance and evolutionary rescue, respectively (based on Fig. 2a). But together they can lead to monopolization effects, which can reduce biodiversity in changing environments.

Although we find that high levels of either dispersal or adaptive potential are sufficient to preserve almost all species in analogue environments (Fig. 2a), we believe that this is unlikely to occur in many natural communities. Empirical estimates suggest that dispersal generally limits species climate ranges over latitudinal gradients, although less so over elevation gradients (45). Furthermore, estimates suggest that rates of climatic niche evolution in vertebrates and many plants are far too slow to keep pace with climate change (31, 32). Instead, we expect that rates will be limiting enough so that dispersal and evolution interactively determine whether species are able to persist under environmental change. Limiting rates of adaptation are especially likely for organisms with long generation times, small populations, and limited dispersal ability, or in fragmented landscapes. Even in bacterial monocultures, where we expect rapid adaptation to occur, evolutionary rescue is facilitated by dispersal (46). Likewise, current evidence for evolutionary rescue in communities of interacting species suggests that both adaptation and dispersal play key roles (25).

Our simulations are simplified representations of how ecological communities respond to ongoing environmental change. However, our findings are qualitatively unchanged by increasing the spatial complexity of our landscapes (Fig. S6) or increasing genetic complexity (Fig. S7). Furthermore, our model also results in rates of phenotypic change (0 to 0.025 haldanes; Fig. S8) that fall within the range estimated in natural populations (47) and within the range of evolutionary rates which can be sustained indefinitely under directional environmental change (34). The degree to which dispersal and adaptive potential have conflicting contributions to the maintenance of biodiversity under environmental change depends on the specific parameters in our model but is pervasive across all the combinations of parameter strengths in our sensitivity analyses (Fig. S10, S11). Of course, additional complexities that are not captured by our model may modify these effects. We suspect that interspecific variability in dispersal and adaptive potential should lead to stronger monopolization effects (22). However, if high variability in rates erodes equalizing coexistence mechanisms (48), pre-change diversity may be reduced, thus precluding strong monopolization effects. Furthermore, we also know that dispersal rates and distance are under selection in changing environments (49) as well as mutation rates (50) and this has the potential to also increase the likelihood of interactions between dispersal and the evolution. Specifically, dispersal is well known to evolve during range shifts (49, 51). However, dispersal can also be highly plastic with respect to environmental gradients (52) and other species (53). While these complexities are beyond the scope of the present study, future theoretical and empirical work will have to account for the eco-evolutionary details of the dispersal process. Despite these open questions, our findings clearly highlight the fact that dispersal and evolution of environmental optima can interact to produce monopolization effects in the community context, and that ignoring this interaction may lead us to overpredict species persistence under future environments.

### Conclusions

Climate change poses a major risk for biodiversity (54) and is already causing the reorganization of ecosystems globally (55). Whether species will keep pace and persist in this changing climate remains uncertain (56) but is expected to depend on both ecological and evolutionary processes (1, 22). Our results clearly highlight dispersal and evolution can interact, creating the potential for monopolization effects, which can result in the loss of biodiversity. Together, our findings provide a more general understanding of the processes that act to maintain biodiversity in a changing world (Fig. 4). This understanding highlights the need for more focus and study on the interactions between ecological and evolutionary processes and how they jointly determine how species and communities will respond to future environmental conditions.

## Methods

We use an individual based model to simulate the temporal dynamics of 80 species in a landscape comprised of 30 habitat patches (see Supplementary Methods for details). To avoid edge effects, patches in the metacommunity are arranged in a ring, with each patch connected to its two adjacent patches (for 2D – 5×30 landscapes see Supplemental Materials Fig. S6). We assume that environmental values increase over the first half of the ring from 0 to *M/2 – 1* in integer steps and decrease following the same rule in the second half. These local conditions are held constant for an initial period of 10,000 generations to allow the simulation to reach quasi-equilibrium. We then gradually increase the environmental conditions in all patches by a total of 5 units over the next 5000 generations. Individuals reproduce sexually and generations are non-overlapping. The reproductive output of each female depends on the match between its phenotype and the local environmental conditions, as well as inter and intraspecific competition with other individuals that are present in the same patch (Supplemental Methods – Equation 1). Individuals are diploid and inherit one randomly chosen allele from each parent. The phenotype is determined by the average of 2 diploid loci (for simulations with 20 loci see Supplemental Materials Fig. S7). During inheritance, new allele values are drawn at a certain rate *m* from a normal distribution with a mean value equal to that of the parent, and a standard deviation equal to a mutation width of *σ*_*mut*_. Thus, we define adaptive potential as *mσ*_*mut*_^*2*^, which is the per allele variation generated in each generation due to mutations. Dispersal is between adjacent patches with a probability *d*. We contrasted dispersal rates (0 to 0.5) and adaptive potential (0 to 3.6 × 10^−4^) that span the range of outcomes from full extinction, to near full persistence in our model. These levels of adaptive potential resulted in rates of phenotypic evolution between 0 and 0.025 haldanes, which falls within the range estimated in natural populations (47). We quantified the proportion of species that persisted over the period of environmental change, distinguishing between regions of the landscape where the final conditions fell within the range present on the landscape (analogue environments) or not (non-analogue environments).

## Acknowledgements

We thank Matthew Osmond, Luc De Meester, Sally Otto, Mary O’Connor and Nathaniel Sharp for valuable discussions and feedback. PLT is supported by Killam and NSERC Postdoctoral fellowships. This is publication ISEM-YYYY-XXX of the Institut des Sciences de l’Evolution – Montpellier.

