## Supplementary Materials for "The conflict between adaptation and dispersal for maintaining biodiversity in changing environments"

**This PDF file includes:**

Supplementary methods
Table S1
Figures S1 to S13
SI References

**Supplementary methods**

*Reproduction, inheritance, and survival*

We assume that reproduction is sexual and that the sex ratio is, on average, 0.5. As individuals are diploid, they inherit one randomly chosen allele from each parent. The phenotype $z_i$ is the average across both alleles. Results from our model are qualitatively the same in simulations with 20 diploid additive loci (Fig. S7). Simulations with this more complex genetic structure are computationally intensive and so we base our main results on the 2 loci model, where we have more replicates. Females choose a mating partner randomly and produce 2 * $\lambda_i$ offspring in order keep λ interpretable at the population level. Offspring are randomly assigned as female or male with equal probability. During inheritance, new allele values are drawn at a certain rate *m* from a normal distribution with a mean value equal to that of the parent, and a standard deviation equal to a mutation width of $\sigma_{mut}$. Thus, we define adaptive potential as $m^*\sigma_{mut}^2$, which quantifies the per allele variation generated in each generation due to mutations.

The reproductive output of each female is determined by a random draw from a Poisson distribution, which simulates demographic stochasticity. After accounting for density regulation (following 1) and environmental mismatch due to local adaptation, the expected value of this distribution $\lambda_i$ for female *i* of species *j* at time *t* in patch *l* is equal to:

$$1)\quad \lambda_{i,j,l}(t) = \frac{\lambda_0}{1+\sum_{k=1}^{S} \alpha_{j,k} N_{k,l}(t)} e^{-\left(\frac{x_l(t)-z_i}{2\sigma_{niche}}\right)^2},$$

where $\lambda_0$ is the fecundity, $\alpha_{j,k}$ is the per capita competitive effect of individuals of species *k* on individuals of species *j*, $N_{k,l}(t)$ is the abundance of species *k* in patch *l* at time *t*, $x_l$(t) - $z_i$ represents the mismatch between the trait $z_i$ and the environment $x_l$(t) experienced by individual *i*, and $\sigma_{niche}$ is the niche width. Thus, reproductive success is unaffected by the phenotype of the males, but males contribute to the success of their offspring through contributing one of their two alleles. See Table S1 for a summary of parameters and tested values. Species interaction parameters were generated such that most, but not all, species could co-exist locally, if they share the same environmental optimum. This was achieved by drawing the strength of intraspecific competition, $\alpha_{j,j}$ from a lognormal distribution with mean 0.002 and variance 0.1, and drawing the strength of interspecific competition, $\alpha_{j,k}$ from a lognormal distribution with mean 0.001 and variance 0.1.

*Dispersal and landscape*

Dispersal is natal and occurs before reproduction. The probability that an individual disperses is governed by a Bernoulli distribution, with a probability equal to $d_i$. Dispersing individuals leave their natal patch and disperse to one of the adjacent patches (nearest-neighbour dispersal) with equal probability.

Mortality during dispersal occurs with probability $\mu$, which summarizes all possible costs of dispersal, be they risk, time or energy costs (2).

Patches in the metacommunity are arranged in a ring, with each patch connected to its two adjacent patches. This ring arrangement allows us to avoid edge effects in our landscape. We assume that environmental values increase over the first half of the ring from 0 to *M/2 - 1* in integer steps and decrease following the same rule in the second half. Results from this simple landscape are qualitatively the same as simulations on a 5x30 patch torus (Fig. S6). But, simulations on this more spatially complex landscape are computationally intensive and so we present results from the simple ring landscape, where more replicates can be run. After a "burn-in" phase of 10,000 generations, environmental change occurs at a constant rate across the landscape such that $x_l(t+1) = x_l(t) + \Delta E$. The magnitude of environmental change was chosen so that 2/3 of the patches would have a pre-change analogue for their environmental conditions at the end of the simulation.

We initialized the landscape by adding individuals (half females, half males) from all species at their single-species equilibrium density $\widehat{N} = {(\lambda_0 - 1)}/{\alpha_{j,j}}$, rounded to the nearest integer, to all patches. Individuals of one species always had the same local adaptation optimum at simulation start, randomly selected from a uniform distribution, that spanned the range of initial environmental conditions.

*Scenarios*
We contrasted 12 dispersal rates *d* spanning the range from 0 to 0.5. We combined this with a factorial comparison of five rates of mutation spanning the range from 0 to 0.1, which result in levels of adaptive potential spanning the range from 0 to 3.6 x $10^{-4}$. These ranges of dispersal and adaptive potential were chosen to cover the range from full extinction to full persistence in our model. These mutation rates are higher than what is typical in natural populations (but see 3). However, these rates were required to allow evolutionary rescue, given the small population sizes that were necessary to make the model computationally feasible and the fact that our model lacks mechanisms for maintaining standing genetic variation. Thus, these mutation rates should not be interpreted as being comparable to those in natural populations, but rather they abstractly represent different levels of genetic variation that affect rates of selection in response to environmental change. Note, increased adaptive potential does not need to depend on increased mutation rates but may also be the consequence of increased mutation widths $\sigma_{mut}$, and our results do not qualitatively change when we vary adaptive potential by varying $\sigma_{mut}$ instead (Fig. S12). For each combination of dispersal and adaptive potential, we ran 50 replicate simulations, each time drawing a new community matrix as described above. We additionally ran a sensitivity analysis for the most relevant parameter values (Fig. S10, S11).

A typical run of the simulation takes 5-10 minutes on a Dell Precision 7910 machine with Intel Xeon E5- 2630 v4 processors and 64 GB RAM. Therefore, generating our main scenario results takes 10 dispersal rates x 5 mutation rates x 100 replicates x 5-10 minutes = 17-34 CPU days. However, these values depend very much on total population sizes and vary between replicates.

*Response variables*
We calculated the proportion of species that were maintained over the course of environmental change at the regional metacommunity scale, separating patches based on whether their final environmental conditions had a pre-change analogue (here, environmental values below 15) or not. In all cases, we compared the communities at the final time step of the burn-in phase ($t$ = 10, 000) with those at the end of the simulation, after environmental change had occurred ($t$ = 15, 000). The proportion of species maintained was calculated as the number of species that were present in the region (analogue or non-analogue) after environmental change, divided by the number of species that were present prior to environmental change. Of course, extinctions can still occur following this period, even without environmental change, but based on simulations where we kept the environment constant for the entire period these were rare and would not have qualitatively affected our findings (Fig. S13).

We calculated the mean change in species range size as the number of habitat patches occupied by a species after change minus the number of habitat patches occupied by that species before change, at the end of the burn-in phase. We included both analogue and non-analogue patches in this calculation, but only included species that persisted over the course of environmental change.

We calculated the interspecific variation in range shift as the standard deviation in range size changes across all species that persisted during environmental change within each replicate run of the model.

We calculated the mean leading vs. trailing range edge expansion as:

$$2)\quad \nu = \frac{\sum_{j=1}^{S_{post}} \left|max\left(x_{j_{post}}\right) - max\left(x_{j_{pre}}\right)\right| - \left|min\left(x_{j_{post}}\right) - min\left(x_{j_{pre}}\right)\right|}{S_{post}},$$

where $x_{j_{pre}}$ and $x_{j_{post}}$ are the environmental conditions in the habitats occupied by species $j$ before and after environmental change, respectively. $S_{post}$ is the regional diversity of the metacommunity, post environmental change. This measure calculates the degree to which the range of each species shifts on its leading edge (min environmental conditions) vs. its trailing edge (max environmental conditions), on average.

**Data accessibility**

The simulation code produced for and used in this study is available on github.
Zenodo DOI:10.5281/zenodo.2019881

**Supplemental Tables**

**Table S1**: Important model parameters, definitions and tested values. Standard values are underlined. A sensitivity analysis including all other values is shown in Figures S2 and S3.

| Parameter | Values | Meaning |
|---|---|---|
| $d$ | 0, 1 x $10^{-5}$, 5 x $10^{-5}$, 1 x $10^{-4}$, 5 x $10^{-4}$, 1 x $10^{-3}$, 5 x $10^{-3}$, 1 x $10^{-2}$, 5 x $10^{-2}$, 1 x $10^{-1}$, 5 x $10^{-1}$ | **dispersal rate**: probability of emigrating for individuals of a given patch |
| $m$ | 0, 0.005, 0.01, 0.03, 0.1 | **mutation rate**: probability of a random change (Gaussian with $\sigma_{mut}$) of a local adaptation allele |
| $\lambda_0$ | 1.5, <u>2</u>, 4 | **fecundity**: maximal mean number of offspring |
| $\mu$ | 0.05, <u>0.1</u>, 0.2 | **dispersal cost**: probability of surviving a dispersal event |
| $\sigma_{niche}$ | 0.5, <u>1</u>, 2 | **niche width**: width of the local adaptation function (Gaussian) |
| $\sigma_{mut}$ | 0.02, 0.04, <u>0.06</u>, 0.08 | **mutation width**: width of the mutation function (Gaussian) |
| $\alpha_{j,j}$ | 0.001, <u>0.002</u>, 0.004 | **mean intraspecific competition coefficient**: intraspecific competition coefficients (i.e., the diagonal of the community matrix) are drawn from a log-normal distribution with mean $\alpha_{j,j}$ and variance 0.1 |
| $\alpha_{j,k}$ | 0.0005, <u>0.001</u>, 0.002 | **mean interspecific competition coefficient**: intraspecific competition coefficients (i.e., off-diagonal elements in a symmetric community matrix) are drawn from a log-normal distribution with mean $\alpha_{ij}$ and variance 0.1 |

### Supplemental Figures

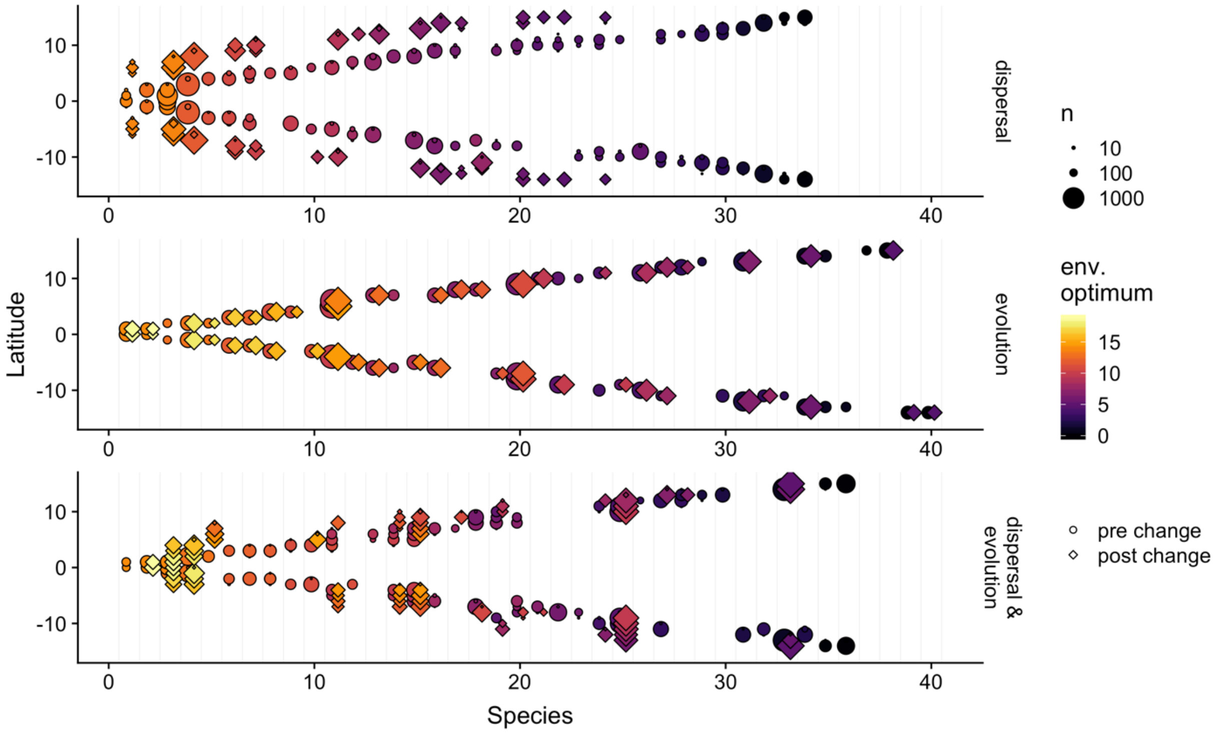


**Figure S1.** Illustration of how dispersal and evolution of environmental optimum, in isolation (a,b, respectively) and in combination (c), affect how species respond to environmental change. Patches on the two sides of the landscape ring are show as positive and negative latitudes, respectively (see Fig. 1 just positive latitudes). Species are arranged on the x-axis by their pre-change latitude. Circles and diamonds are paired for each species and indicate the latitude and size of each population prior to and after environmental change, respectively. Species with circles but not diamonds failed to persist. The colour shows the mean environmental optimum in each population. Panel a shows a scenario where dispersal is intermediate (0.01) and the adaptive potential is zero; here, species respond to environmental change by shifting to higher latitudes to maintain the match between their phenotype and their local environmental conditions. Panel b shows a scenario where dispersal is 0 and adaptive potential is intermediate ($1.08 \times 10^{-4}$); here species respond to environmental change through adaptation (change in colour), with no change in latitude. Panel c shows a scenario where dispersal (0.01) and adaptive potential ($1.08 \times 10^{-4}$) are both intermediate; here species respond through a combination of shifting latitude and through adaptation. Results shown are from one representative simulation run with standard parameter values (Table S1).

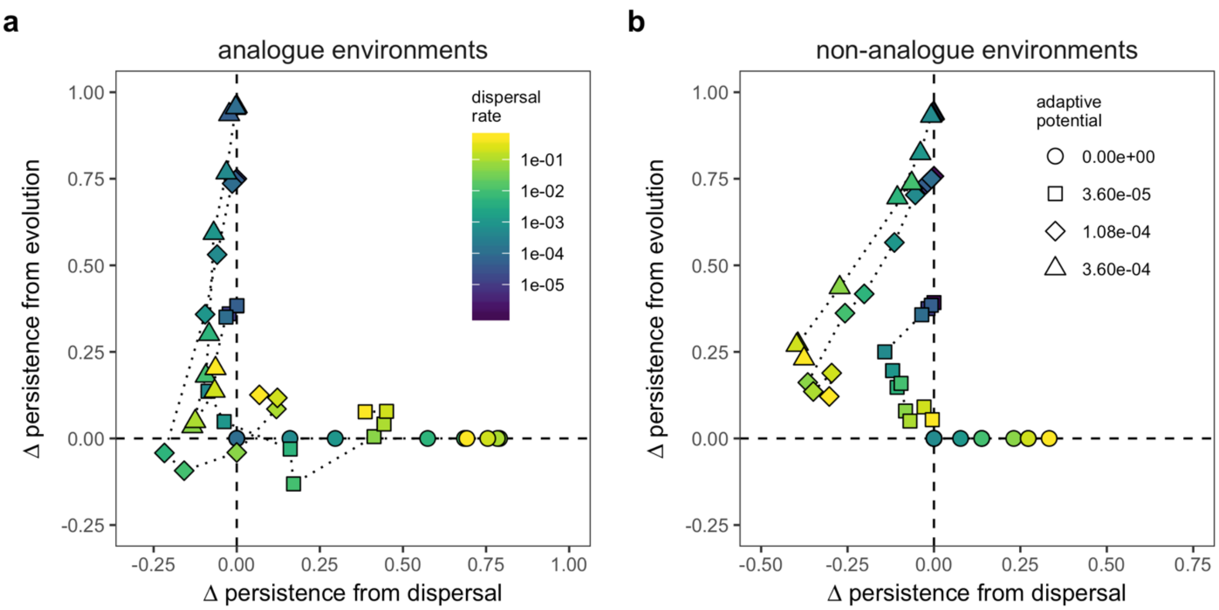


**Figure S2.** The proportion of species maintained in each combination of dispersal and adaptive potential compared to if dispersal were 0 (x-axis) or if adaptive potential was 0 (y-axis) in analogue (a) and non-analogue environments (b). This indicates the degree to which the combination of dispersal and adaptive potential alters the persistence of species compared to if dispersal or evolution were to operate in isolation. Points that fall below zero on the x-axis indicate cases where dispersal reduces the number of species that could persist with that level of adaptive potential in the absence of dispersal. The y-axis shows the same information but for adaptive potential. The dispersal rates and adaptive potential values are shown by the colour and shape of the points, respectively. The dotted lines connect points with the same adaptive potential in increasing dispersal rate. Median values across all 50 replicates with standard parameter values (Table S1) are shown.

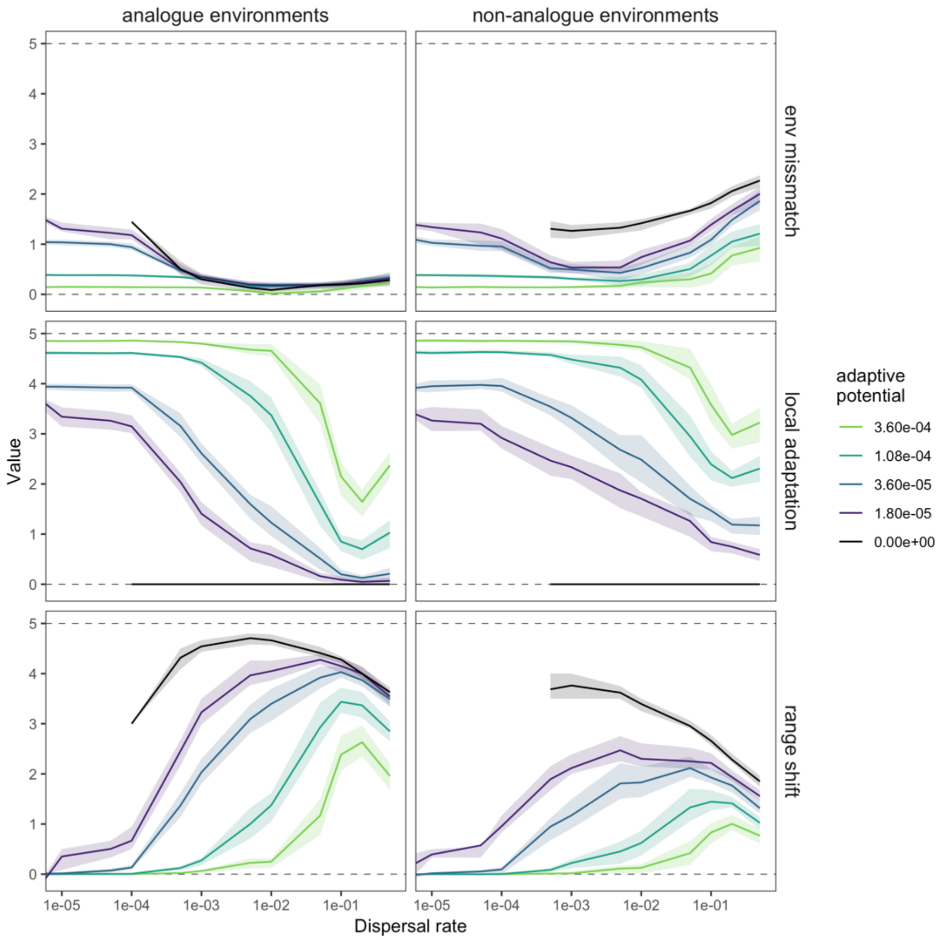


**Figure S3.** The degree of mismatch between individuals and their local environmental conditions at the end of the simulations (top row), the average magnitude of trait change that surviving lineages have undergone over the course of environmental change (middle row), and the magnitude of the range shift that surviving lineages have undergone (bottom row) depending on dispersal and adaptive potential (colour). We separate these results based on whether the final environmental conditions in each patch fall within the pre-environmental change range of conditions (analogue; left column) or not (non-analogue; right column). The lines show the median value across 50 replicate simulations with standard parameter values (Table S1) and the bands show the interquartile range.

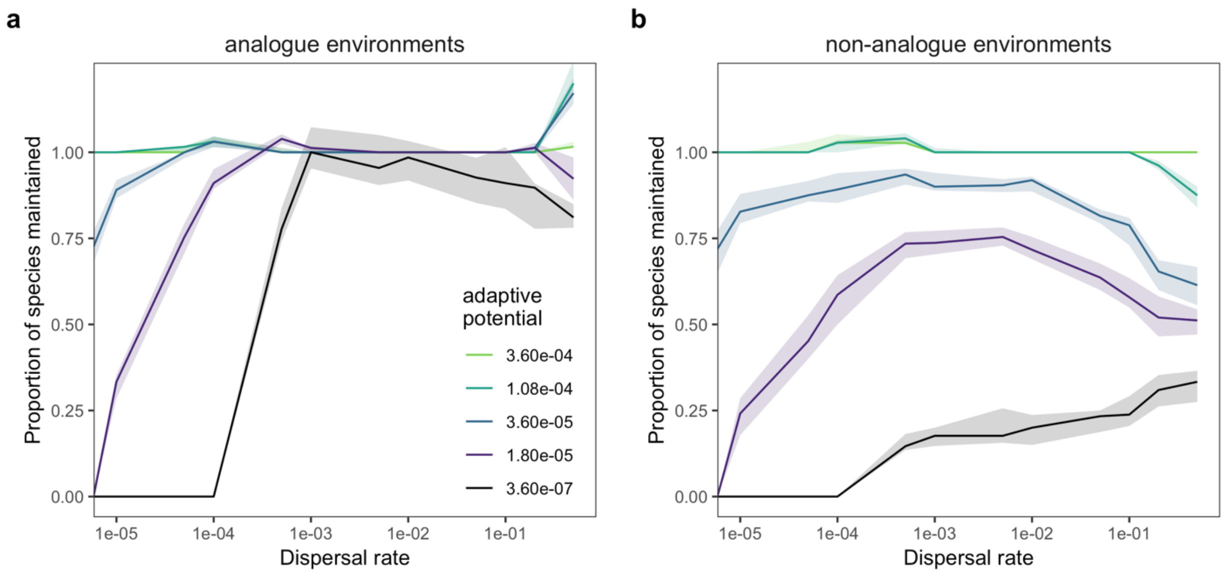


**Figure S4.** The proportion of species that are maintained following environmental change depending on dispersal and adaptive potential (colour) when all interspecific competition $\alpha_{j,k}$ is set to 0. We separate these results based on whether the final environmental conditions in each patch fall within the pre-environmental change range of conditions (a, analogue) or not (b, non-analogue). The lines show the median value across 25 replicate simulations with standard parameter values (Table S1) and the bands show the interquartile range. Note, this simulation was run with an extremely low adaptive potential of 3.6 x $10^{-7}$ instead of 0.

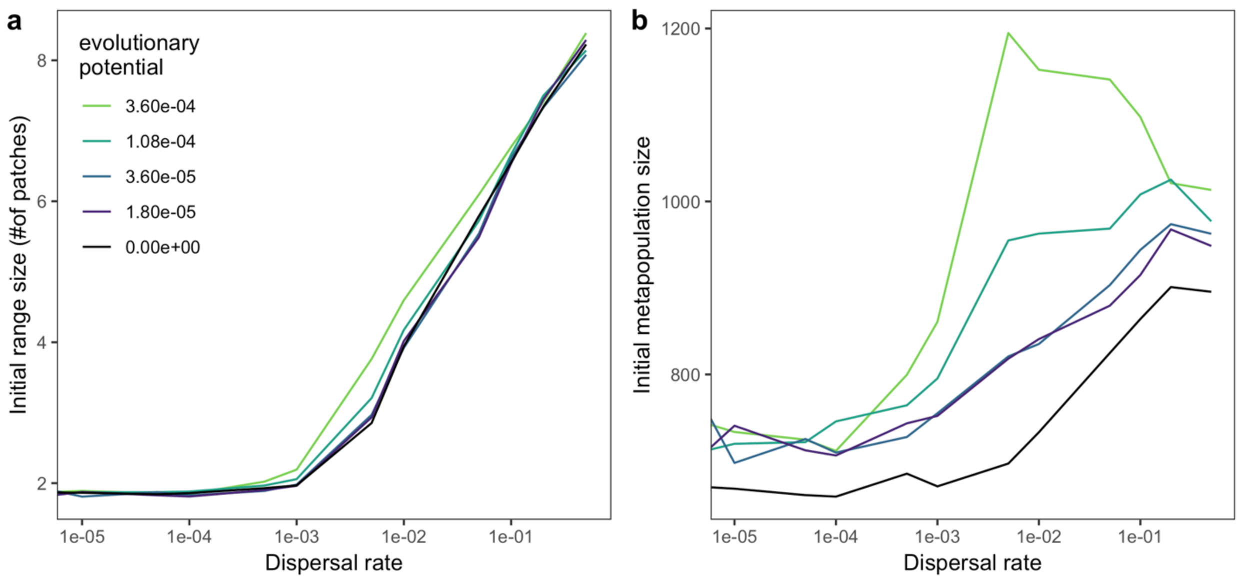


**Figure S5.** The average range size (a) and total metapopulation size (b) of species prior to environmental change, depending on dispersal and adaptive potential (color). Metapopulation size is the average number of individuals of each species across all patches. The lines show the median value across 50 replicate simulations with standard parameter values (Table S1).

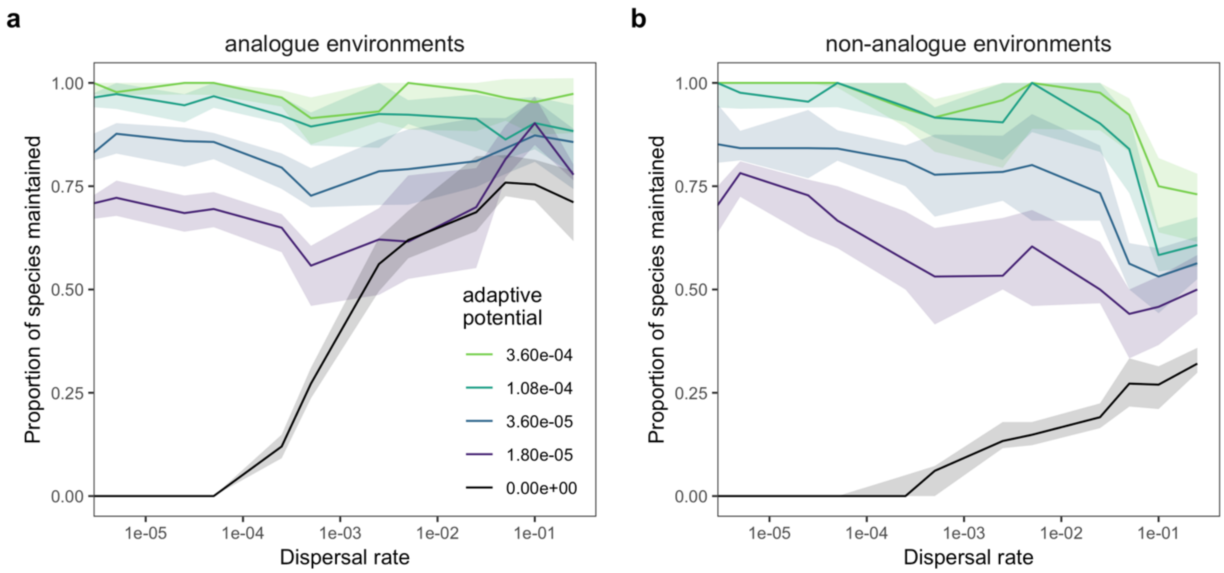


**Figure S6.** Sensitivity analysis showing the proportion of species maintained following environmental change in a 5x30 patch torus. Just like in the simple ring landscapes, the environment increases over the front half of the torus and decreases over the back half of the torus to avoid edge effects. More species persist in this landscape compared to the ring landscape with a given level of adaptive potential. This is because overall metapopulation sizes are larger (x5) and so favourable mutations are more likely. The lines show the median value across 20 replicate simulations.

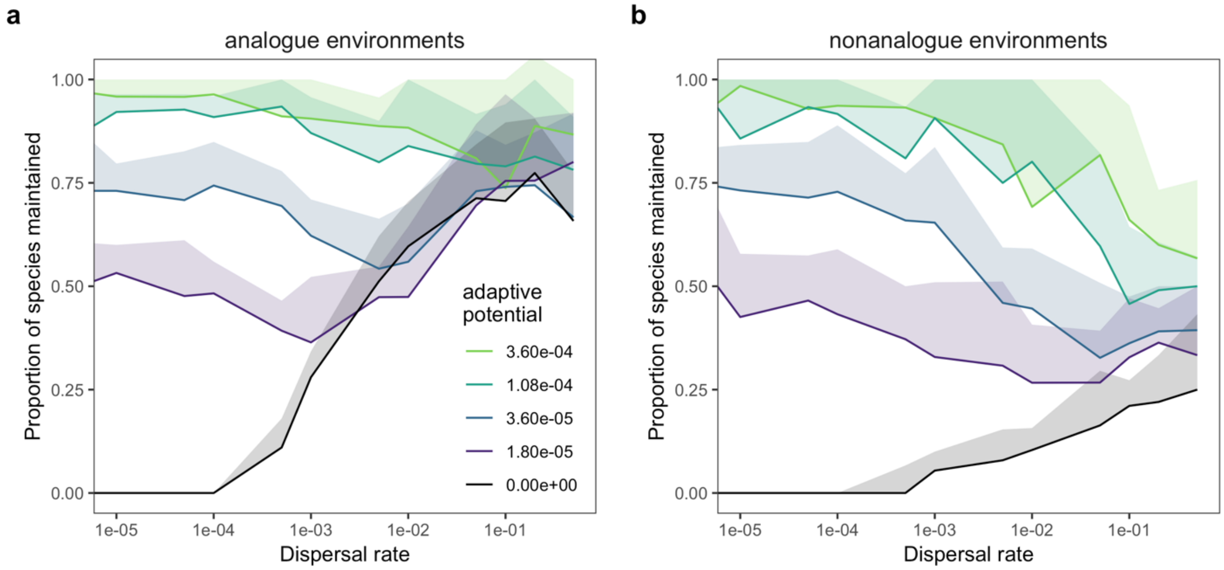


**Figure S7.** Sensitivity analysis showing the proportion of species maintained following environmental change in a model where the phenotype is determined by 20 diploid additive loci (see also 4). The simulations are identical to the standard setting with the difference that every individual is characterized by 20 diploid loci that additively determine the phenotype. $\sigma_{mut}$ was lowered to take the different genetic structure into account ($\sigma_{mut}$ = 0.0125). The lines show the median value across 20 replicate simulations.

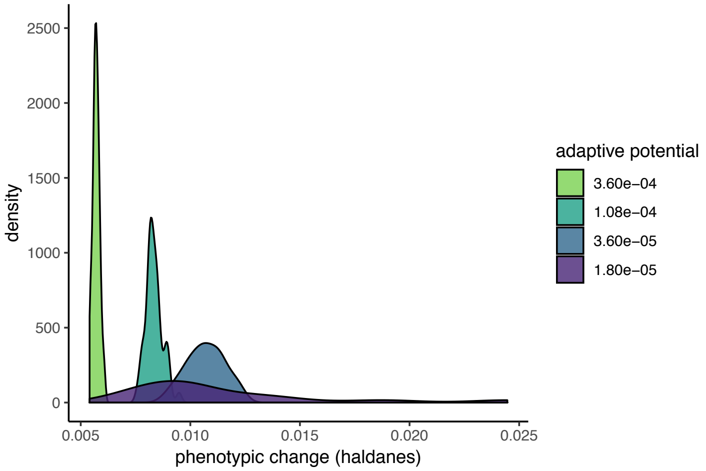


**Figure S8. R**ates of phenotypic change over the period of environmental change in surviving populations in the no dispersal scenario expressed as standard deviations per generation (haldanes) following Hendry and Kinnison (5). We only compare populations that persist in the same habitat patches; thus, we exclude rates for simulations with dispersal because dispersal results in range shifts. Note, that high adaptive potential results in lower haldanes, because phenotypic change is standardized by phenotype variation, and high adaptive potential results in greater variation in phenotype. Thus, the amount of absolute phenotypic change required for species to persist decreases with adaptive potential. The large variation in haldanes seen at the lowest adaptive potential is due to the fact that very few populations are able to persist with this level of adaptive potential, but the ones that do vary greatly in their absolute phenotypic change and the amount of within population variation in their phenotypes. These estimate rates of phenotypic change fall well within the range reported from natural populations (5). However, estimates from natural populations are generally from relatively few generations and it is likely that rates of phenotypic change would be slower over long periods of environmental change. Still our rates of phenotypic change are less that those that can be sustained indefinitely under directional environmental change (6).

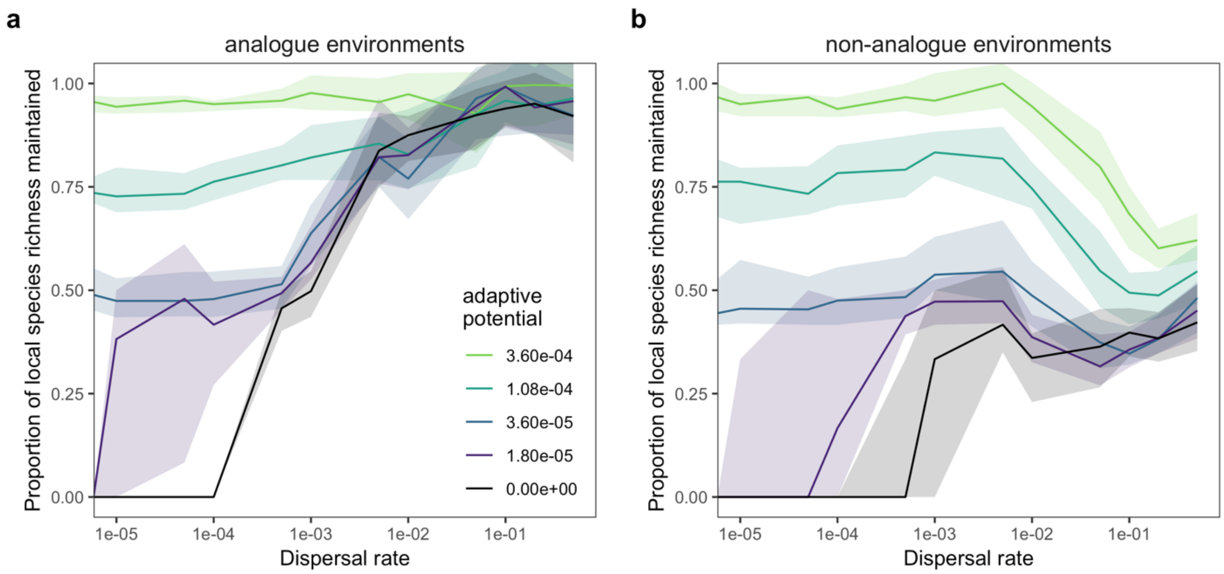


**Figure S9.** The proportion of species that are maintained in each patch (local scale richness) following environmental change depending on dispersal and adaptive potential (colour). The proportion of species maintained was calculated as the number of species that were present in each patch (a - analogue or b - non-analogue) after environmental change, divided by the number of species that were present before. The lines show the median value across 50 replicate simulations with standard parameter values (Table S1) and the bands show the interquartile range.

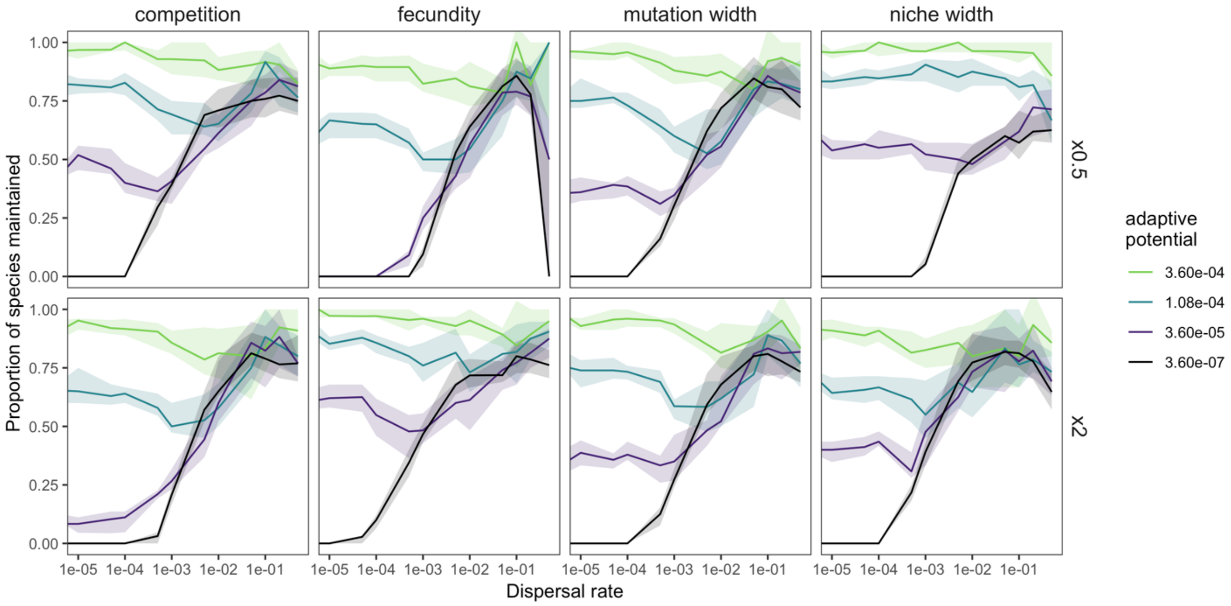


**Figure S10.** Sensitivity analysis showing the proportion of species maintained in analogue environments when fixed parameters are halved (top row) or doubled (bottom row). Each column shows the results when a single parameter is varied, keeping all of the others standard as outlined in Table S1. These are, from left to right, competition, fecundity, mutation width, and niche width. One exception is that the reduction in fecundity is x0.75 (1.5 vs 2). The lines show the median value across 25 replicate simulations with standard parameter values (Table S1) and the bands show the interquartile range. Note, these simulations were run with an extremely low adaptive potential of $3.6 \times 10^{-7}$ instead of 0.

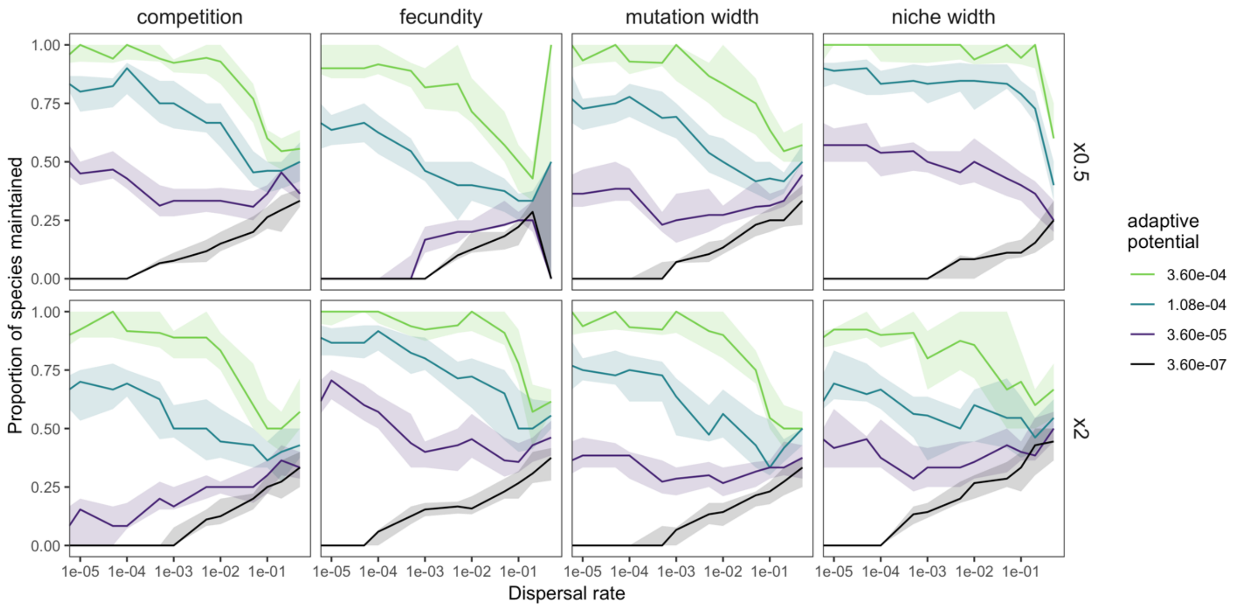


**Figure S11.** Sensitivity analysis showing the proportion of species maintained in non-analogue environments when fixed parameters are halved (top row) or doubled (bottom row). Each column shows the results when a single parameter is varied, keeping all of the others standard as outlined in Table S1. These are, from left to right, competition, fecundity, mutation width, and niche width. One exception is that the reduction in fecundity is x0.75 (1.5 vs 2). The lines show the median value across 25 replicate simulations with standard parameter values (Table S1) and the bands show the interquartile range. Note, these simulations were run with an extremely low adaptive potential of $3.6 \times 10^{-7}$ instead of 0.

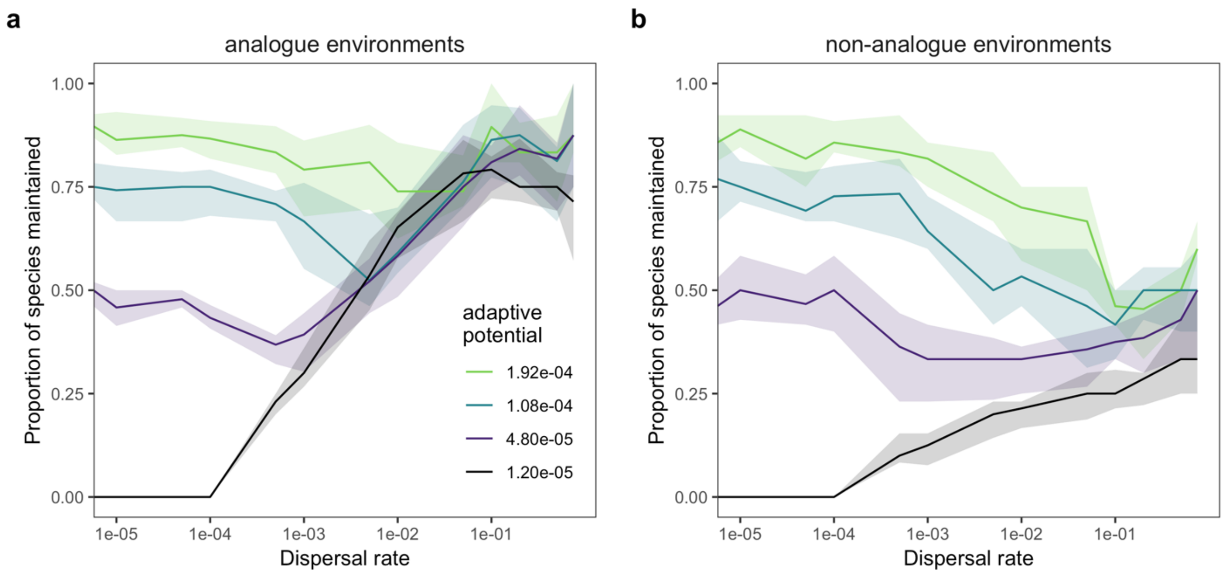


**Figure S12.** The proportion of species that are maintained in each patch (local scale richness) following environmental change depending on dispersal and adaptive potential (colour). Here adaptive potential is varied by contrasting different mutation widths $\sigma_{mut}$ (0.02, 0.04, 0.06, and 0.08) with a constant mutation rate *m* of 0.03. The proportion of species maintained was calculated as the number of species that were present in each patch (a - analogue or b - non-analogue) after environmental change, divided by the number of species that were present before. The lines show the median value across 50 replicate simulations with standard parameter values (Table S1) and the bands show the interquartile range.

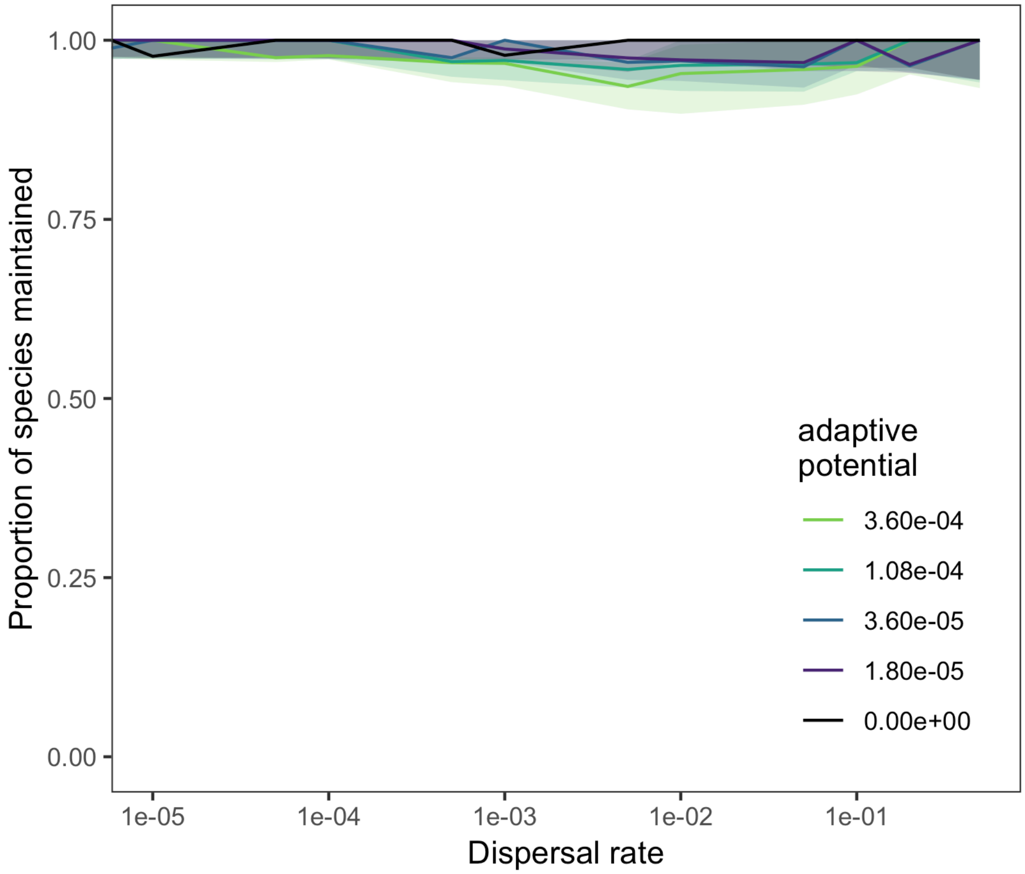


**Figure S13.** The proportion of species that are maintained across all habitat patches between time steps 10,000 and 15,000 in the absence of environmental change. The lines show the median value across 50 replicate simulations with standard parameter values (Table S1) and the bands show the interquartile range.
